# Directed evolution of an orthogonal transcription engine for programmable gene expression in eukaryotes

**DOI:** 10.1101/2024.09.25.614978

**Authors:** Shaunak Kar, Elizabeth C. Gardner, Kamyab Javanmardi, Daniel R. Boutz, Raghav Shroff, Andrew P. Horton, Thomas H. Segall-Shapiro, Andrew D. Ellington, Jimmy Gollihar

## Abstract

T7 RNA polymerase has enabled orthogonal control of gene expression and recombinant protein production across diverse prokaryotic host chassis organisms for decades. However, the absence of 5’ methyl guanosine caps on T7 RNAP derived transcripts has severely limited its utility and widespread adoption in eukaryotic systems. To address this shortcoming, we evolved a fusion enzyme combining T7 RNAP with the single subunit capping enzyme from African swine fever virus using *Saccharomyces cerevisiae*. We isolated highly active variants of this fusion enzyme, which exhibited roughly two orders of magnitude higher protein expression compared to the wild-type enzyme. We demonstrate the programmable control of gene expression using T7 RNAP-based genetic circuits in yeast and validate enhanced performance of these engineered variants in mammalian cells. This study presents a robust, orthogonal gene regulatory system applicable across diverse eukaryotic hosts, enhancing the versatility and efficiency of synthetic biology applications.

## Introduction

For decades, the single-subunit phage RNA polymerase T7 RNAP has been a cornerstone in biotechnology applications including recombinant protein production ^1,2^. T7 RNAP is particularly useful due to its high specificity to its small 17 bp promoter, which enables high yield RNA production while operating independently of the native transcriptional machinery ^3–5^. In addition to its routine use in protein overexpression, the orthogonal, predictable qualities of T7 RNAP have led to its use in complex genetic circuitry. In particular, T7 RNAP and its engineered derivatives have been programmed into controllers ^6–8^ resource allocators ^9^, autoregulatory circuits ^3,10^, and Boolean logic programs^11^.

While T7 RNAP systems have predominantly been utilized in *Escherichia coli*, T7 RNAP has also been adapted for recombinant protein production in a variety of prokaryotic hosts of industrial importance including *Pseudomonas*, *Bacillus*, and *Corynebacterium*^10,12,13^. In contrast, porting T7 RNAP driven protein expression systems into eukaryotes has posed challenges. Unlike prokaryotes, functional eukaryotic mRNAs undergo significant post-transcriptional modifications critical for mRNA translation, nuclear export, and stability^14,15^. These modifications include the addition of a 5’ methyl guanosine cap (5’cap) and stretches of adenosine residues (polyA) in the 3’ untranslated region (UTR). Eukaryotic mRNA processing involves multiple coordinated steps synchronized with RNA polymerase II initiation, elongation, and termination^16–18^. The absence of these modifications in T7 RNAP-derived transcripts has limited the adaptation of T7 RNAP for protein expression in eukaryotes^19^.

Among these modifications, polyA addition can be encoded in specific DNA sequences and appropriately processed post-transcription^20^. However, adding the 5’cap to T7-derived transcripts has remained challenging^19–21^. Previous attempts to recruit host capping machinery by fusing Pol II-derived signaling domains to T7 RNAP were unsuccessful in appreciable protein expression^22^. However, it has been shown that T7-expressed transcripts can be modified using viral capping enzymes (CEs), which can catalyze all steps in cap formation independently of the host capping machinery ^23–26^. Among these, the vaccinia capping enzyme (VCE) from the vaccinia virus plays a crucial role in cytoplasmic capping and translation of virus-encoded genes necessary for replication^27–30^. Co-expression of T7 RNAP and VCE can produce capped T7-derived mRNAs and subsequent protein translation in the cytoplasm of human cells without relying on host transcriptional and capping machinery ^25,26,31^. However, the levels of capped transcripts are low relative to the uncapped transcripts^31^, possibly due to its uncoordinated activity in an otherwise well controlled multi-step pathway.

Recently, it has been shown that fusion enzymes composed of T7 RNAP and the VCE could mediate protein expression in the cytosol of mammalian cells^23^, and that this fusion enzyme showed enhanced protein expression compared to the unlinked enzymes^23^. Additionally, alternative capping enzymes were evaluated as fusion partners, with the capping enzyme from African swine fever virus (NP868R) showing the highest protein expression in HEK293T cells^23^. Despite these advances, T7-fusion mediated protein expression was reported to be much lower than native nuclear expression of reporter proteins, indicating that there is potential for enhancing the enzyme’s functionality ^23^. Additionally, while a theoretical advantage of the T7-fusion enzyme is broad-host compatibility in eukaryotes, its performance has not yet been benchmarked in a non-mammalian background.

In this study, we develop a yeast-based directed evolution platform to engineer and study the activity of the fusion enzyme comprised of NP868R and T7 RNAP to generate functional mRNA transcripts and achieve protein expression in a cap-dependent manner. We found multiple variants with enhanced co-transcriptional capping activity of the fusion enzyme. We isolated highly active variants that demonstrate nearly two orders of magnitude higher activity than the wild-type fusion enzyme. Using these highly active variants, we demonstrate that gene expression can be predictably controlled using previously characterized T7 RNAP-based genetic elements, greatly expanding the repertoire of T7 RNAP-based genetic circuitry in eukaryotes, and show enhanced activity in mammalian cell lines.

## Results

### Activity of NPT7 fusion in yeast is low

We investigated the activity of a fusion enzyme composed of NP868R and T7 RNAP (referred to as NPT7) in *Saccharomyces cerevisiae*, as previous studies have only characterized its function in mammalian cells for cytoplasmic expression, not for nuclear synthetic circuitry ^23,32^. Using yeast as a eukaryotic model, we sought to test NPT7’s ability to generate capped transcripts of target genes in the nucleus under the control of the T7 RNAP promoter (**Fig. 1a**). Given the importance of the 5’ cap for efficient translation in yeast, we hypothesized that protein production levels would directly reflect the activity of both NP868R and T7 RNAP. We constructed a gene encoding NPT7, fused with a glycine-serine linker and an N-terminal nuclear localization signal (NLS), and placed it under the control of a galactose-responsive promoter (pGal) ^33^. This cassette was integrated into the HO locus of the yeast genome ^33^ (**Fig. 1a**). Additionally, we designed a reporter construct where a fluorescent protein, ZsGreen, was controlled by the T7 RNAP promoter, followed by a polyadenylation signal and a T7 terminator encoded in a multi-copy yeast plasmid ^33^ (**Fig. 1a**). Upon transforming these constructs into *Saccharomyces cerevisiae* strain BY4741 and galactose induction, we observed a two-fold increase in protein production (**Fig. 1b,c**). To determine the specific role of the capping enzyme, we introduced a point mutation in NP868R that disrupts cap formation, generating capping ‘dead’ or null versions of NPT7 ^32^ (**Fig. S1a**). Transformation and induction of these mutant enzymes resulted in negligible reporter expression (**Fig. S1b**), confirming the critical role of the capping activity by the wildtype fusion enzyme.

**Figure 1:**
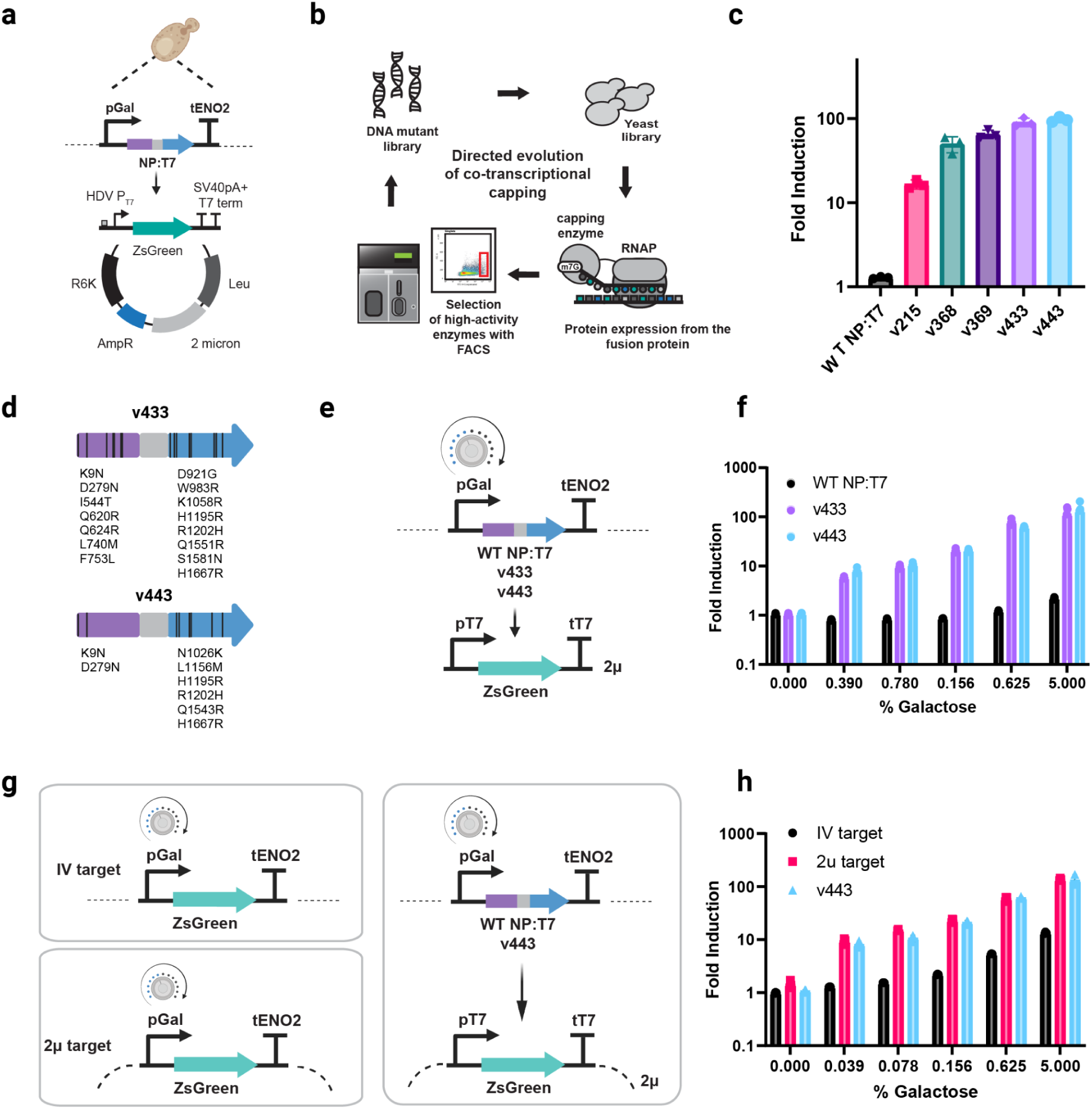
**A)** Design of NPT7 expression cassette under the control of the Gal promoter integrated in the HO locus along with a reporter plasmid consisting of ZsGreen under the control of T7 RNAP promoter followed by a polyadenylation signal on a 2 micron plasmid with an RK6 origin of replication. Upstream of the T7 promoter is an HDV ribozyme insulator. **B)** Directed evolution schematic to generate variants of NPT7 for improved contranscriptional capping activity. **C)** Characterization of evolved variants of NPT7 relative to wild-type after rounds of selection. Fold induction of reporter expression following galactose induction. **D)** List of mutations in v433 and v443 observed in the NP868R and T7 RNAP. **E,F)** Characterization of reporter expression using graded galactose induction for WT NPT7, v433 and v443. **G,H)** Reporter expression under the control of the Gal promoter assayed both integrated in the genome as well as encoded as a multi-copy yeast plasmid

### Directed evolution of NPT7 improves function in yeast

Next, we hypothesized that the activity of the NPT7 fusion enzyme could be enhanced through directed evolution. We designed a selection scheme using error-prone PCR to generate a library of NPT7 variants in yeast, followed by fluorescence activated cell sorting (FACS) to isolate top-performing variants based on fluorescent reporter expression (**Fig. 1b**). After several rounds of selection, we isolated two variants, v433 and v443, which showed nearly two orders of magnitude increase in signal relative to the wildtype (**Fig. 1c**). Genotyping revealed 15 amino acid mutations in the v433 (seven in the NP868R domain and eight in T7) and ten in the v443 (two in NP868R and six in T7) (**Fig 1d**, **Table S1**). Between the two evolved variants, we observed convergence at five residues (K9N, D279N, H1195R, R1202H, and H1667R). Capping-null variants of each variant [33] confirmed that the increased activity primarily relied on capping function, where capping-null variants showed several fold lower reporter expression than functional versions (**Fig. S1b**). Capping-null enzymes of evolved variants showed elevated reporter activity compared to the inactivated version of the WT NPT7, suggesting a low level of background capping activity in the nucleus. Notably, the evolved NPT7 did not show significant activity on genomic targets in yeastwhich might be due to repressive chromatin conditions. Therefore, all testing was performed on high copy yeast plasmids.

### Tunable control synthetic circuits is maintained across kingdoms

We further investigated whether we could tune reporter expression levels by controlling the amount of fusion enzyme expressed. Using a ΔGal2 strain to achieve titratable levels of expression based on galactose concentration^33^, we demonstrated that reporter expression was proportional to the amount of fusion enzyme expressed (**Fig. 1f**). Comparison of reporter expression by v433 and v443 against host-regulated promoters showed that the level of expression from v443 was comparable to the endogenous Gal promoter (**Fig. 1h**).

To determine the nuclear activity of the evolved variants, the genes were cloned without the NLS tags. In *S. cerevisiae* it is known that the reporter plasmid is localized in the nucleus, and thus the removal of the NLS should lead to decrease in the expressed reporter levels (**Fig S2a**). Upon transformation and induction of the non NLS versions of enzymes we see a significant reduction in the expressed levels of reporter compared to the NLS versions thus suggesting that activity of the enzymes is indeed nuclear (**Fig S2b**).

Furthermore, to simplify circuit design and minimize the use of repeated sequences, we sought to characterize the activity of the evolved enzyme variants using different endogenous yeast polyadenyation signals^33^. We systematically swapped the SV40 pA signal sequence to three different yeast signals – tSsa1, tAdh1 and tTdh1 ^33^(**Fig S3a**). For both the evolved variants, we observe that the reporter expression was similar to all the pA sequences thus suggesting that these sequences can be systematically swapped.

One significant advantage of T7 RNAP-driven gene expression is the ability to tune transcriptional response using well-characterized T7 RNAP promoter elements ^3,7–9^ ^34^. In particular we chose to characterize a set of promoter variants with decreasing strength determined both *in vitro* as well as in *E.coli* ^3,35^ in a eukaryotic context using the highly active v433 and v443 variants (**Table S2**). Swapping ZsGreen with other fluorescent proteins (BFP and mScarlet-I) demonstrated that the increased activity was not protein-specific (**Fig. 2a, 2b, S4a**). We also tested three promoter variants known to provide graded levels of attenuated transcriptional activity, leading to proportional protein expression. The rank order of promoter variants was conserved across all circuit designs, indicating predictable control over gene expression orthogonal to host machinery (**Fig. 2c, 2d, S4b**).

**Figure 2:**
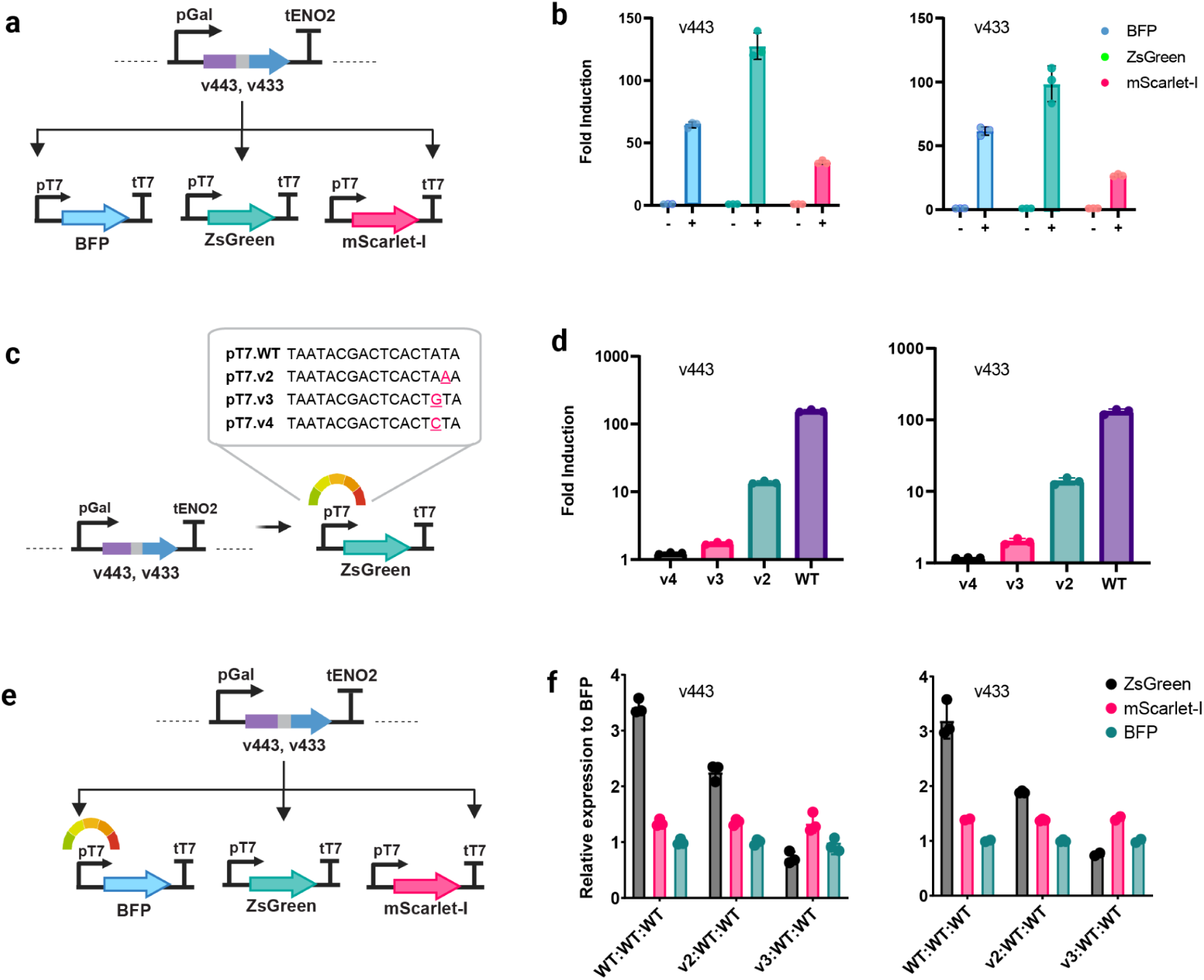
**A)** Reporter constructs encoding fluorescent proteins Zsgreen, BFP, and mScarlet-I were individually transformed into strains containing the v433 and v443 expression cassettes. **B)** Characterization of all reporter expression for both v433 and v443 strains. **C)** List of mutant T7 RNAP promoters to enable tunable gene expression under the control of v433 and v443. **D)** Reporter expression from mutant T7 RNAP promoters shows a graded response consistent with the predicted strength of the promoters. **E)** Design of reporter construct for multiplexed gene expression where only the promoter driving ZsGreen is systematically changed where the other fluorescent cargos are the under the control of the WT T7 RNAP promoter. F) Relative expression of each fluorescent cargo relative to BFP for both v433 and v443 strains.

### Demonstration of multiplexed & orthogonal gene expression circuits by NPT7 variants

To test the capabilities of T7 RNAP to mediate tunable control of multiple targets in parallel, we designed a genetic circuit composed of three separate cargos for multiplexed gene expression using multiple promoter elements simultaneously (**Fig 2e**). Each fluorescent protein was controlled by its own T7 promoter, while only the promoter driving ZsGreen expression was systematically varied with the expression-level modulating mutant versions as described previously (**Fig 2e**). Normalized gene expression of each protein indicated that only ZsGreen levels varied according to the promoter strength, while mScarlet-I and BFP levels remained unchanged, showing independent control of gene expression levels from a single, master regulator enzyme orthogonal to host transcriptional regulation (**Fig 2f**).

Previously, T7 RNAP variants have been engineered to recognize distinct mutant T7 promoters with minimal cross reactivity ^6^. This panel of orthogonal T7 RNAPs may be useful for engineering high complexity pathways independent of host regulation, however these variants have never been ported into eukaryotic hosts. To adapt these tools for protein expression in eukaryotes, we created five orthogonal variants of v443 by grafting specific DNA binding domain mutations. Each variant was tested against its cognate promoter driving a target gene. The results showed successful switching of promoter specificity and capped transcript generation, although protein production levels were lower than the WT version, suggesting the need for further optimization (**Fig. 3a, 3b**).

**Figure 3:**
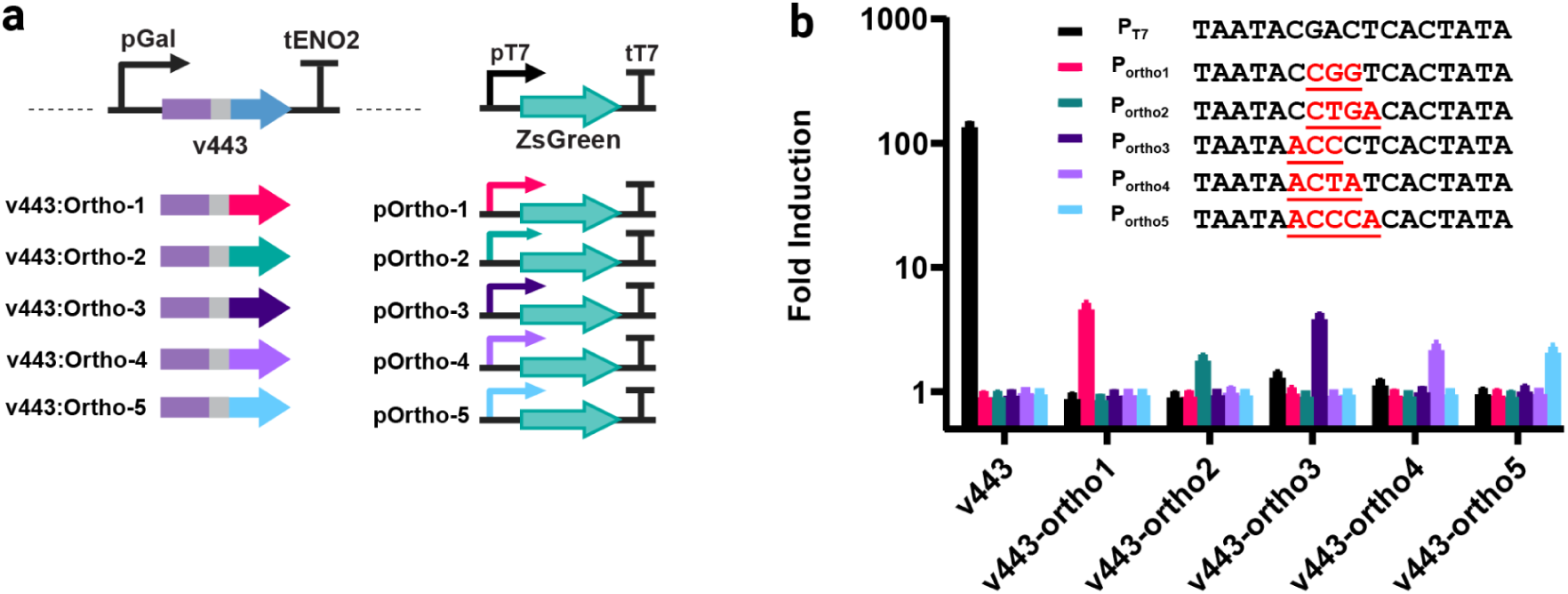
**A)** Five orthogonal variants of v443 constructed by grafting the specific DNA binding domains of T7 RNAP previously characterized. All variants were cloned under the Gal promoter and integrated into the HO locus. A panel of 5 reporter constructs designed with different T7 RNAP promoters driving reporter expression. **B)** Characterization of all six v443 variants activity against the panel of six reporter constructs.

### Evolved NPT7 variants enhance circuit performance in mammalian cells

Given the conserved nature of 5’ cap recognition and translation in eukaryotes, we hypothesized that our engineered variants could function across different eukaryotic chassis with improved activity. To validate this, we characterized the cytoplasmic activity of v433 and v443 in HEK293T cells. These variants, cloned without their NLS and controlled by the CAG promoter and the beta-globulin polyadenylation signal^36^, showed 3-4 fold higher reporter expression compared to the WT in two different reporter contexts comprised of different 5’ and 3’ untranslated regions (UTRs) previously used in the expression SARS CoV2 mRNA vaccine candidates ^37,38^ (**Fig. 4a, 4b, 4c**). This indicates that despite being evolved in yeast, the engineered variants demonstrate enhanced performance across diverse eukaryotic systems, suggesting broad applicability.

**Figure 4:**
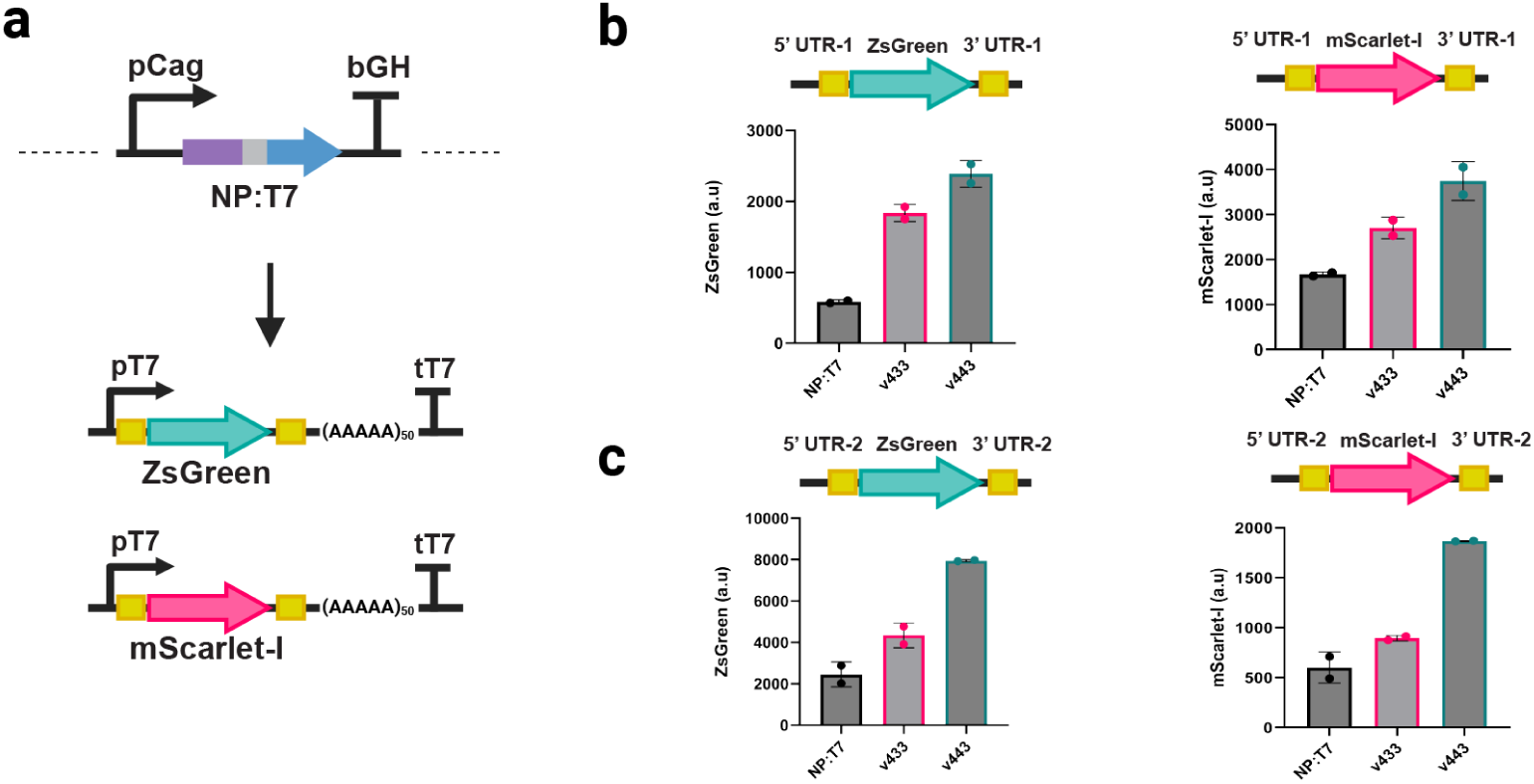
**A)** WT NPT7, v433 and v443 were cloned under the control of CAG promoter followed by polyadenylation signal after the removal of the NLS. Two different reporter construct backbones were designed that included unique 5’ UTR and 3’UTR sequences. In each of the backbone two different fluorescent proteins were used - ZsGreen and mScarlet-I. **B, C)** Activity of WT NPT7, v433 and v443 were characterized in HEK293T cells after transfection with each reporter plasmid.

## Discussion

The development of a fully orthogonal transcription engine capable of expressing 5′ capped mRNAs independently of host RNA polymerase (RNAP) and associated factors represents a significant advancement in the field of synthetic biology. Currently, orthogonal control of gene expression in eukaryotes primarily relies on synthetic promoters regulated by artificial transcription factors ^39–42^. Although substantial progress has been made in designing and characterizing these transcription factors and constructing complex genetic circuits, their dependence on host RNAP dynamics inherently limits their robustness and portability across diverse host systems ^43^. In this study, we demonstrated that an orthogonal RNAP, specifically a fusion of T7 RNAP with the viral capping enzyme NP868R, can efficiently generate functional mRNAs in *Saccharomyces cerevisiae* without relying on the host’s native RNAP and associated factors. However, the initial utility of such fusion enzymes in constructing complex genetic circuits was constrained by their relatively low expression capacity. To address this limitation, we employed a directed evolution strategy, successfully yielding variants with markedly enhanced activity. These evolved variants not only increased protein expression by nearly two orders of magnitude compared to the wild-type fusion enzyme but also maintained orthogonality and functionality across different eukaryotic systems.

Our results demonstrate that these evolved variants enable the construction of genetic circuits with predictable and tunable behavior using well-characterized genetic elements. We observed that the rank order of gene expression driven by T7 promoters was conserved across kingdoms, underscoring the predictability and robustness of our system. Furthermore, the enhanced performance of these variants in HEK293T mammalian cells suggests that our directed evolution approach effectively produces variants with elevated activity that remain largely independent of host-specific factors and interactions. This broadens the applicability of our approach, making it suitable for a wide range of eukaryotic systems beyond yeast.

One of the major challenges in synthetic eukaryotic circuit design, particularly in complex non-model organisms, is the epigenetic silencing of transgene expression^43^. By decoupling transcription and capping of target genes from host factors, our fusion enzymes offer the potential to engineer target genes that are resilient to epigenetic modifications. Specifically, these variants could be further evolved to function effectively under repressive chromatin conditions across diverse eukaryotic hosts. Such robust variants would facilitate the bottom-up construction of genetic circuits resistant to epigenetic silencing, thereby enhancing the stability and predictability of synthetic gene networks in various eukaryotic organisms. Moreover, while capping-T7 can continue to be optimized for improved activity in yeast, challenges such as genomic integration under restrictive chromatin configurations remain. However, opportunities exist to express the fusion enzyme in the cytosol using systems like OrthoRep ^44^, leveraging yeast’s robustness as a chassis for engineered metabolic pathways producing high-value compounds ^45–48^. We envision that capping-T7 could be instrumental in refactoring these pathways for greater stability and fine-grained flux tuning, thereby enhancing metabolic engineering efforts.

Although the performance of our capping-T7 variants is currently superior in yeast compared to mammalian cells, future engineering endeavors may further optimize their expression and activity in mammalian systems. Engineered capping-T7 enzymes hold promise for long-term overexpression in mammalian backgrounds with enhanced resistance to epigenetic silencing, facilitating the recombinant expression of proteins, including biologics, with improved stability and yield. Capping-T7 possesses significant potential across many *in vitro* applications, particularly in the synthesis of 5′ capped mRNAs, which are foundational to RNA therapeutics such as vaccines. T7 RNAP is the primary enzyme employed in these applications and has been extensively engineered to optimize its utility ^49,50^. Our evolved capping-T7 enzymes could simplify the manufacturing process by enabling single-step synthesis of capped RNA, thereby streamlining production workflows. While each application may require project-specific optimizations, our work demonstrates the evolvability of capping-T7, its portability across kingdoms, and provides a robust template for developing orthogonal gene expression circuitries in eukaryotes.

---

**Figure S1:**
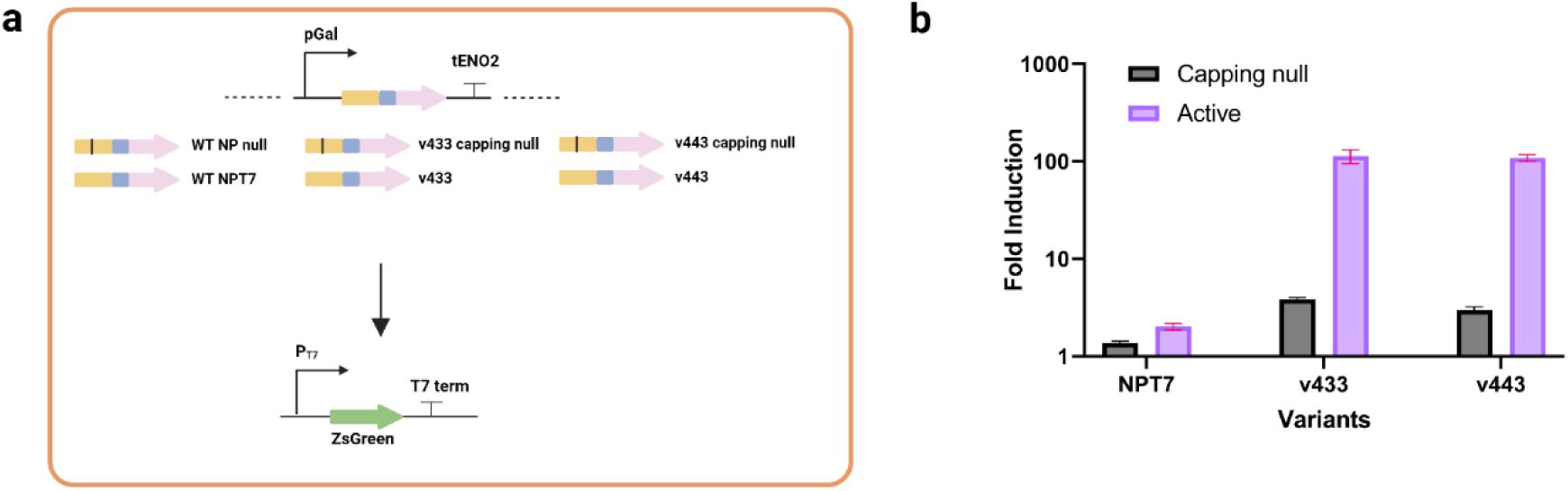
A) Design of the capping null variants of WT NPT7, v433 and v443 by introducing K282A mutation that has been previously shown to abrogate capping activity. B) Characterization of reporter expression for each capping null variant.

**Figure S2:**
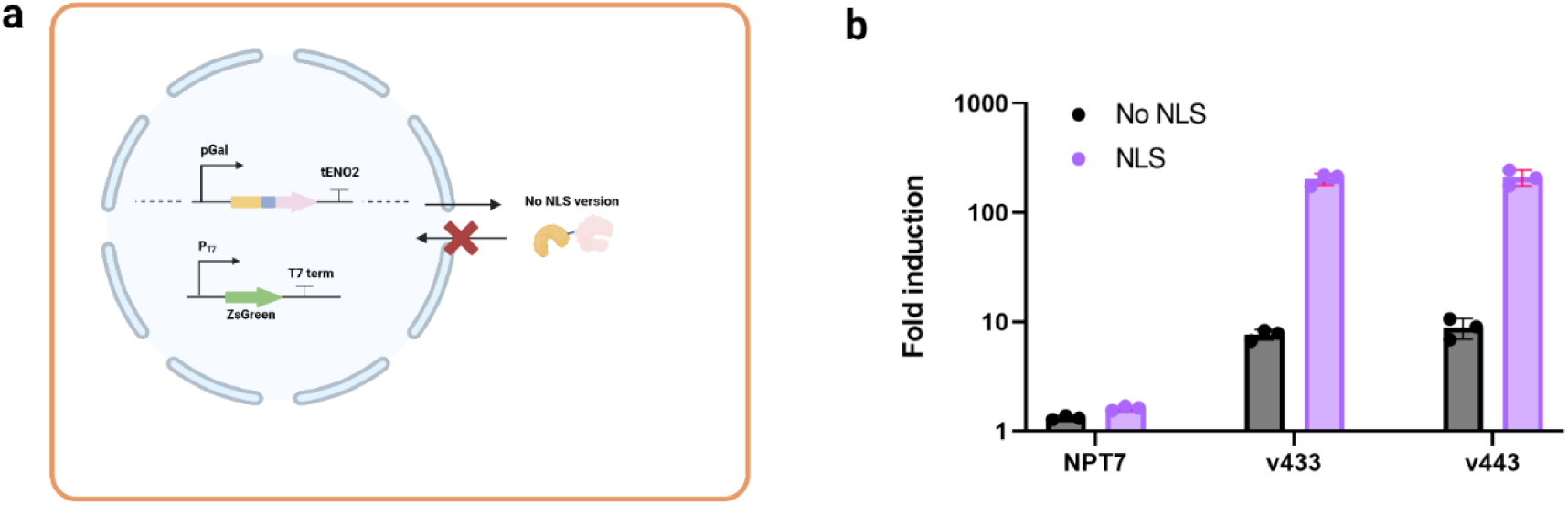
Absence of a nuclear localization signal (NLS) leads to reduction in expressed reporter gene. **A**) Design of constructs without the NLS and transformed into strains containing the reporter construct. **B**) Characterization of the reporter expression for both NLS and without NLS versions of the WT NPT7, v433 and v443 respectively.

**Figure S3:**
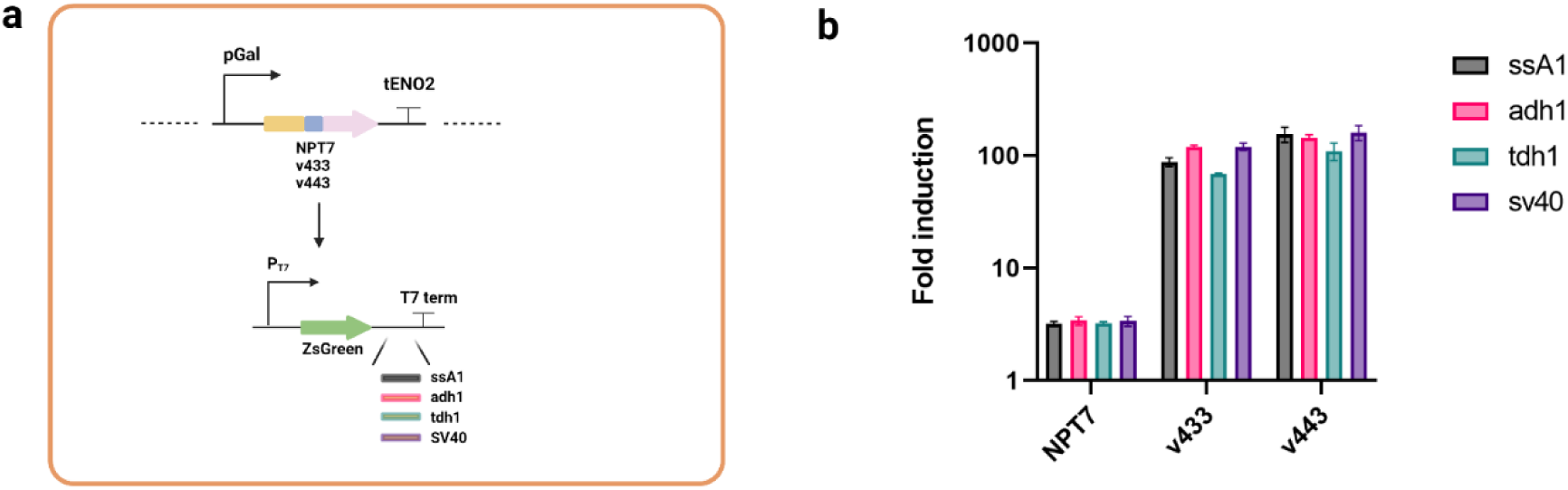
Use of yeast different polyadenylation signals for reporter expression. **A)** Three different yeast polyadenylation signals were used – tSsa1, tAdh1, tTdh1 along with SV40. **B)** The reporter expression was characterized for the different constructs under both the evolved variants as well as the wild-type version

**Figure S4:**
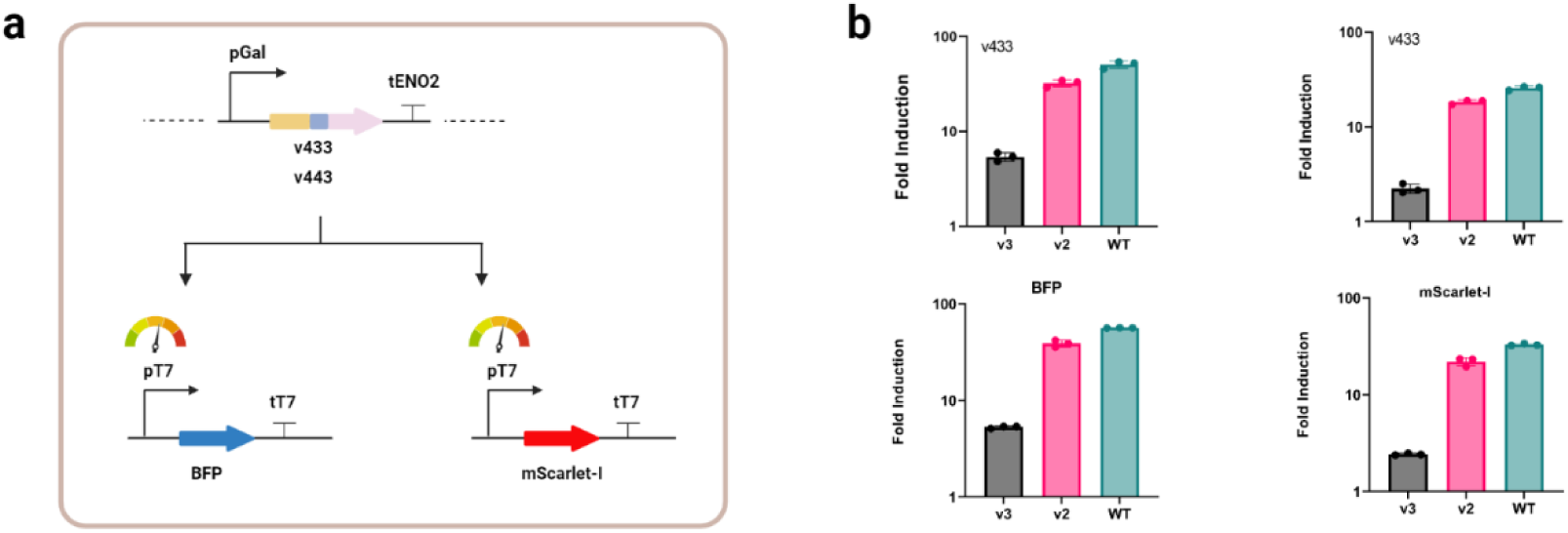
**A)** Mutant T7 RNAP promoter variants v2 and v3 controlling the expression of two fluorescent cargos - BFP and mScarlet-I. **B**) Characterization of each reporter expression under the control of the evolved variants - v433 and v443.

**Table S1:**
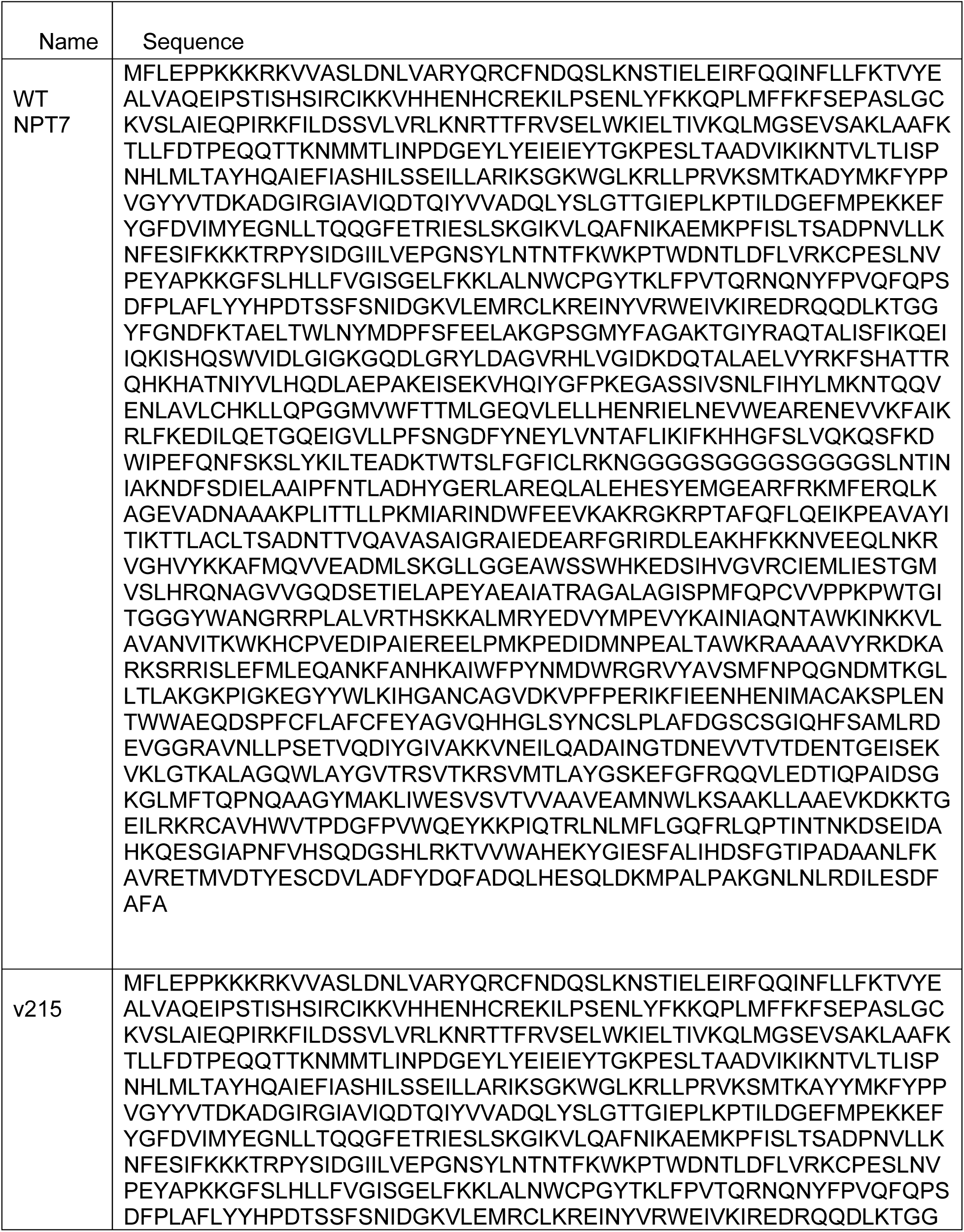
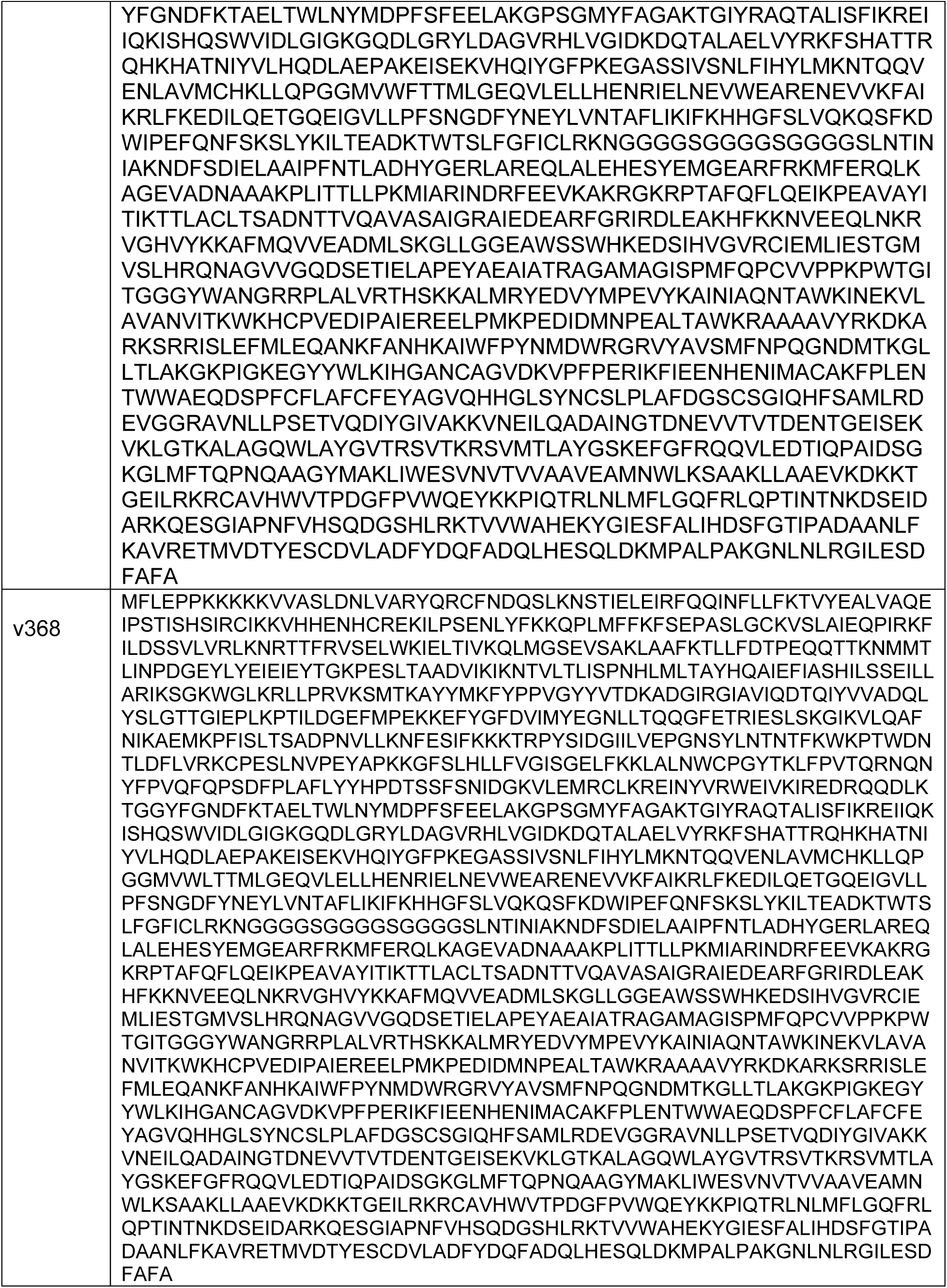
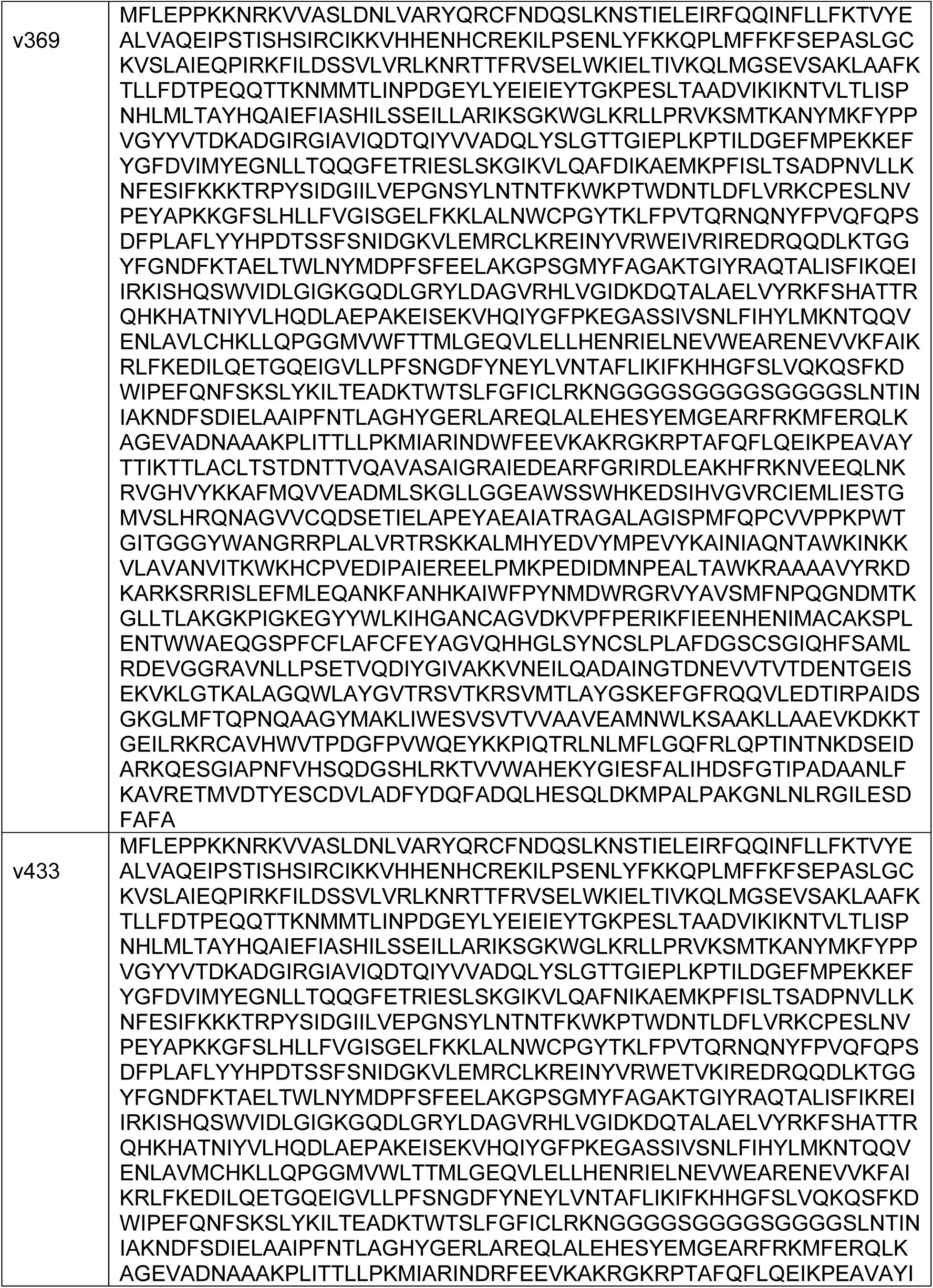
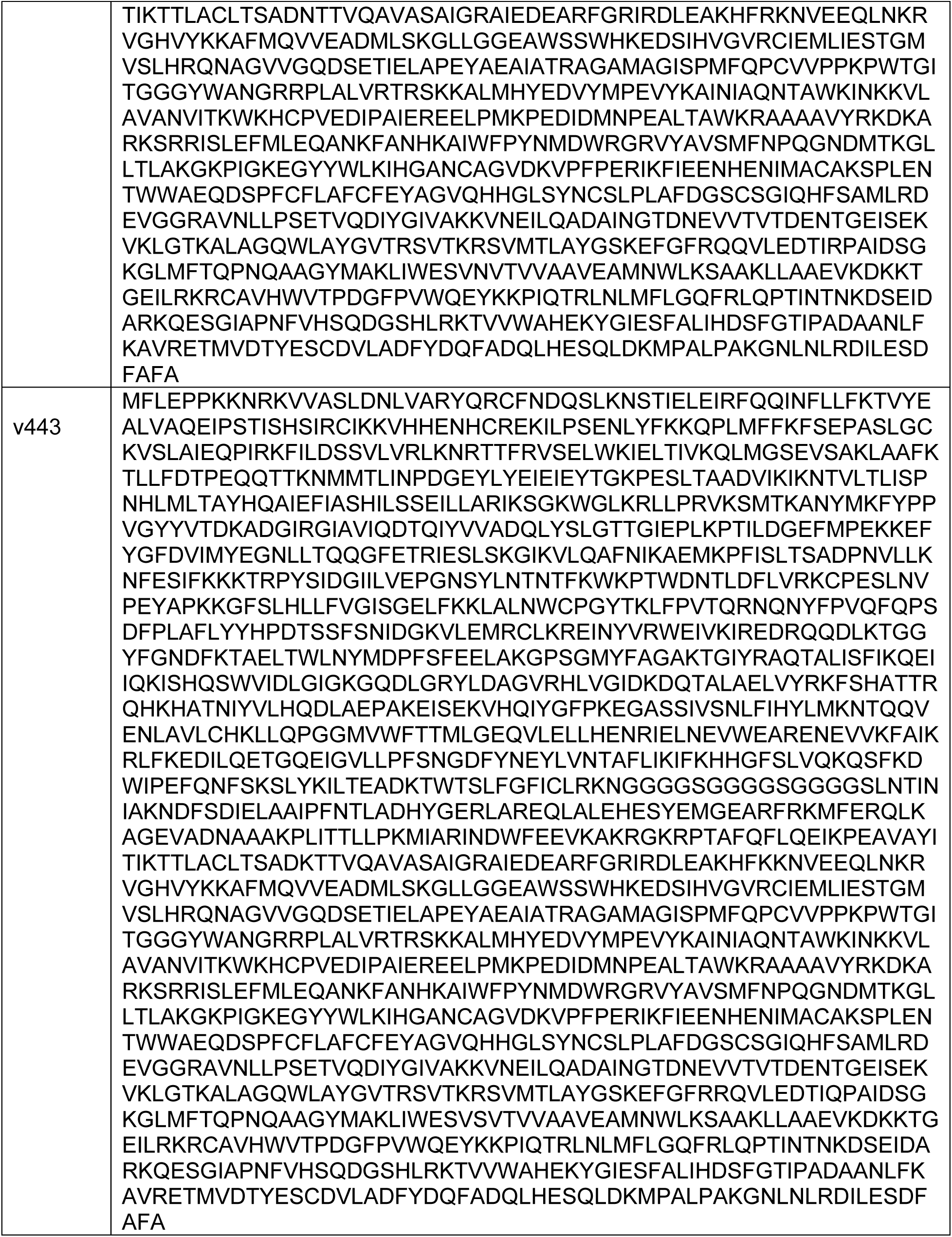
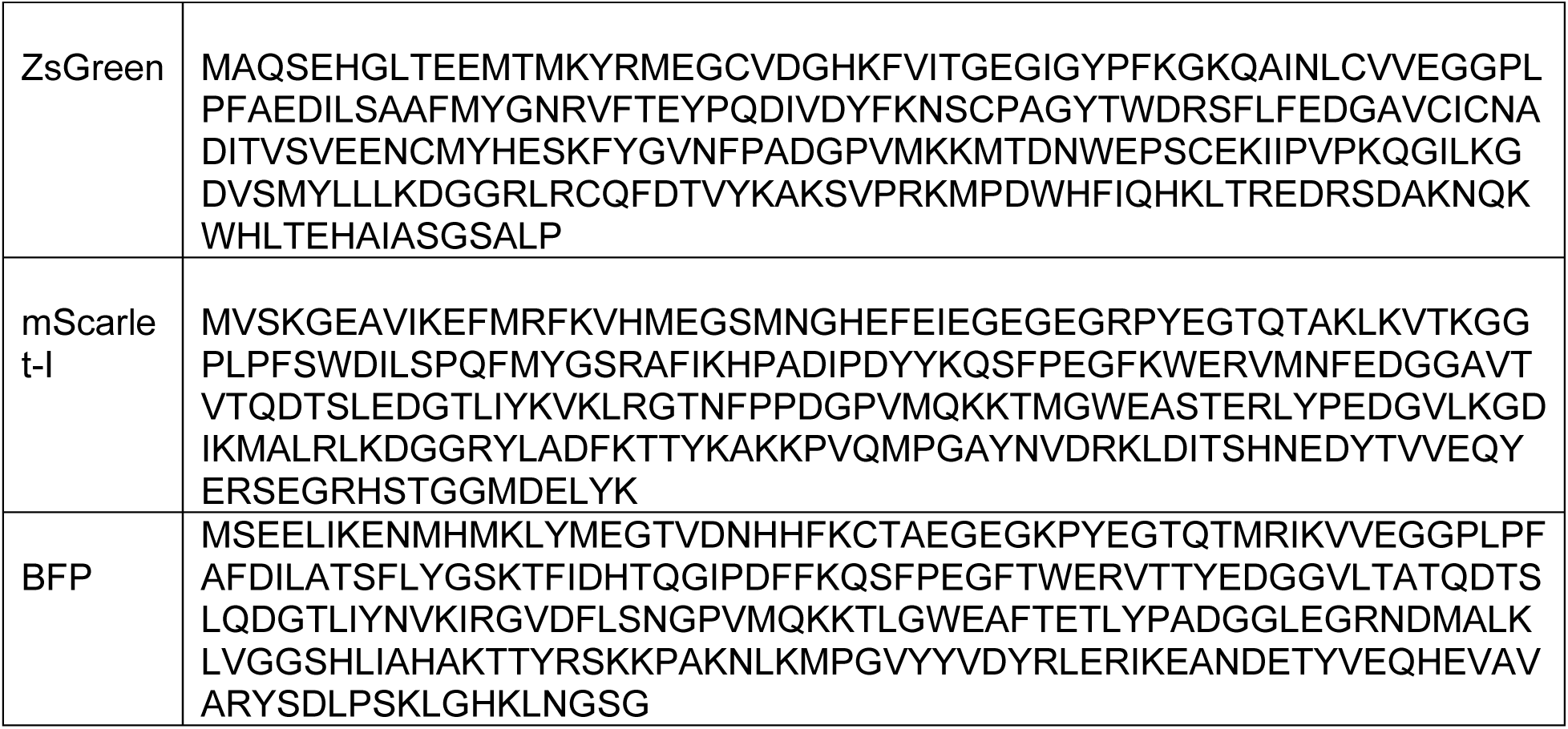
List of coding sequences.

**Table S2:**
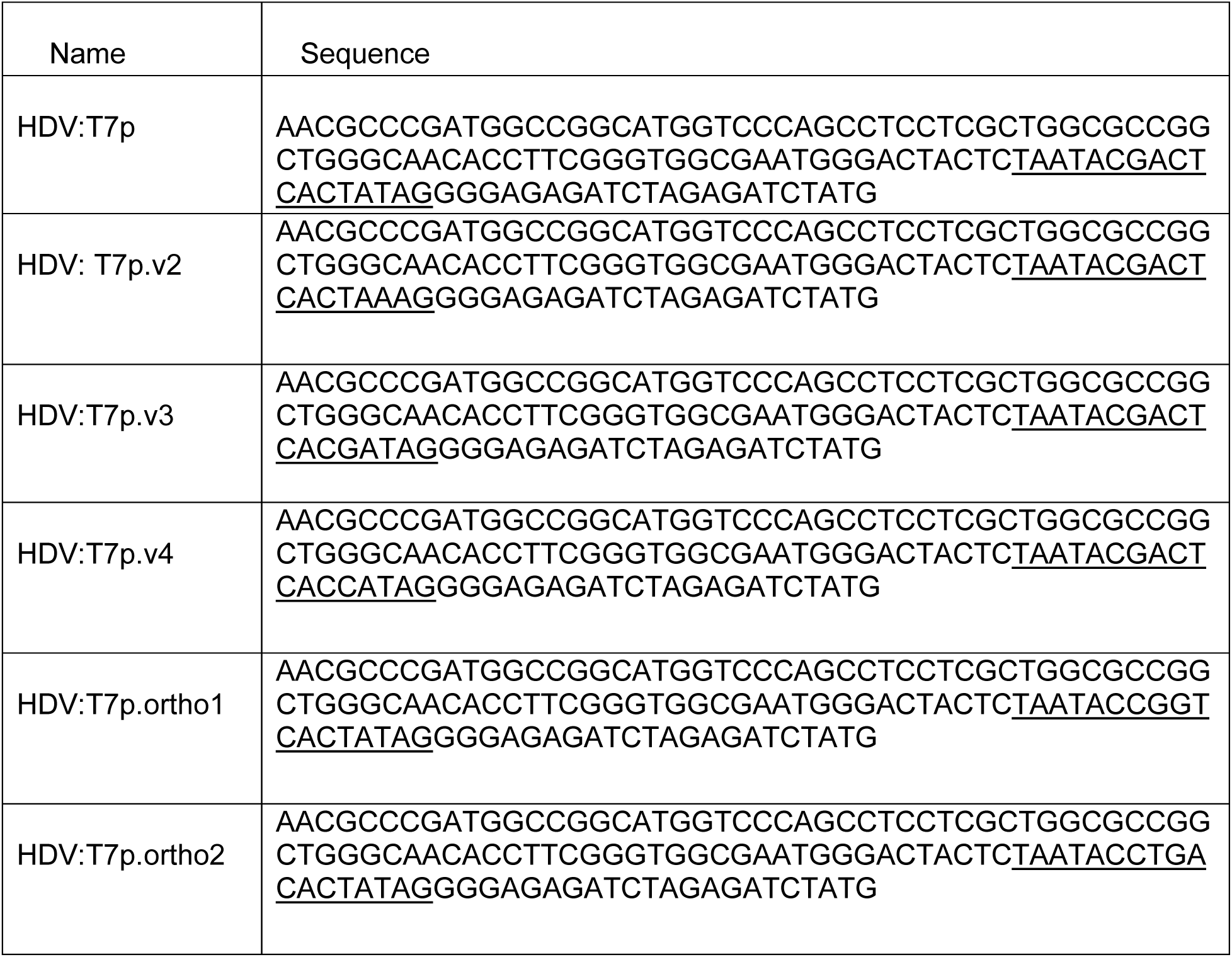
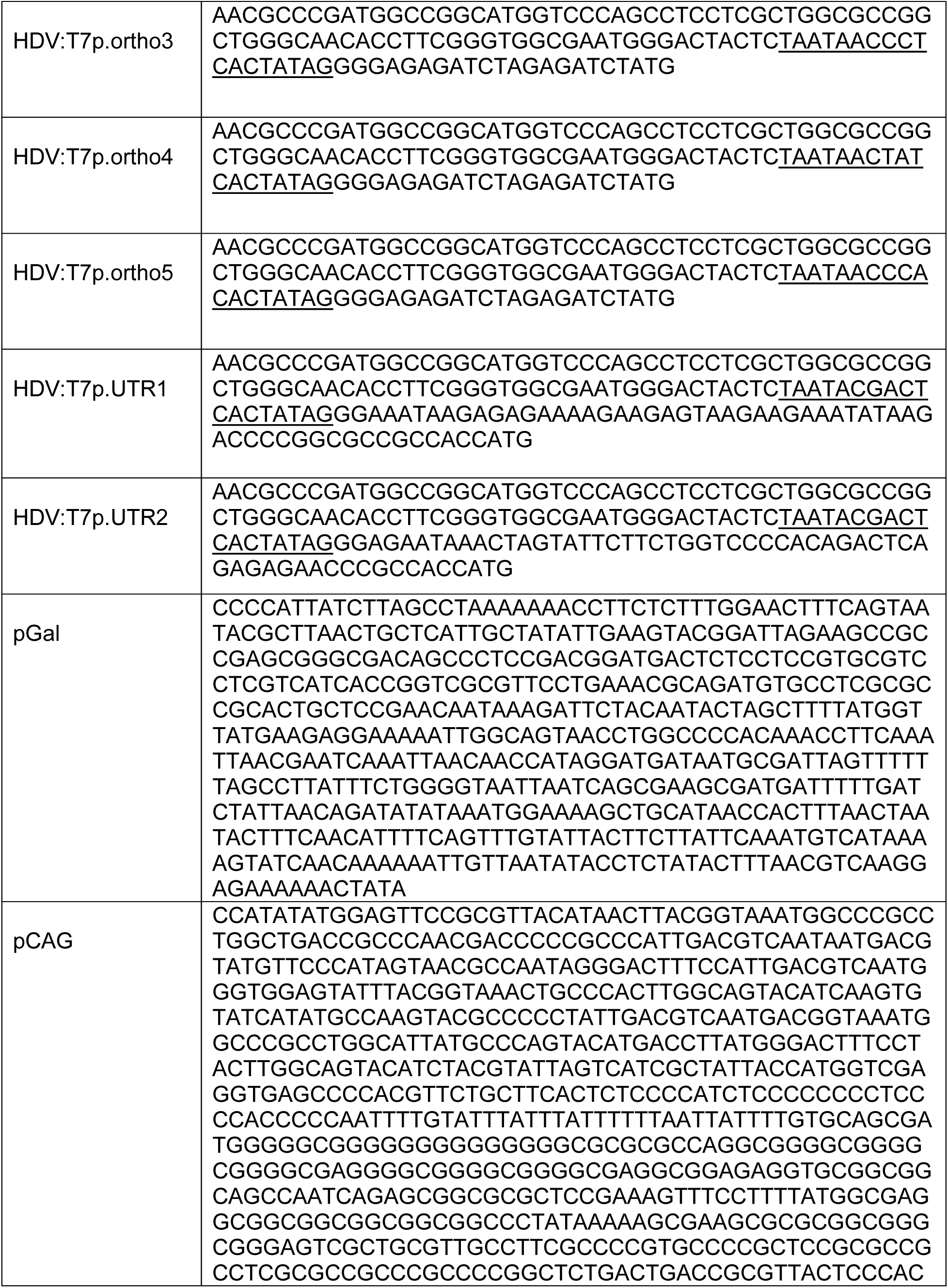
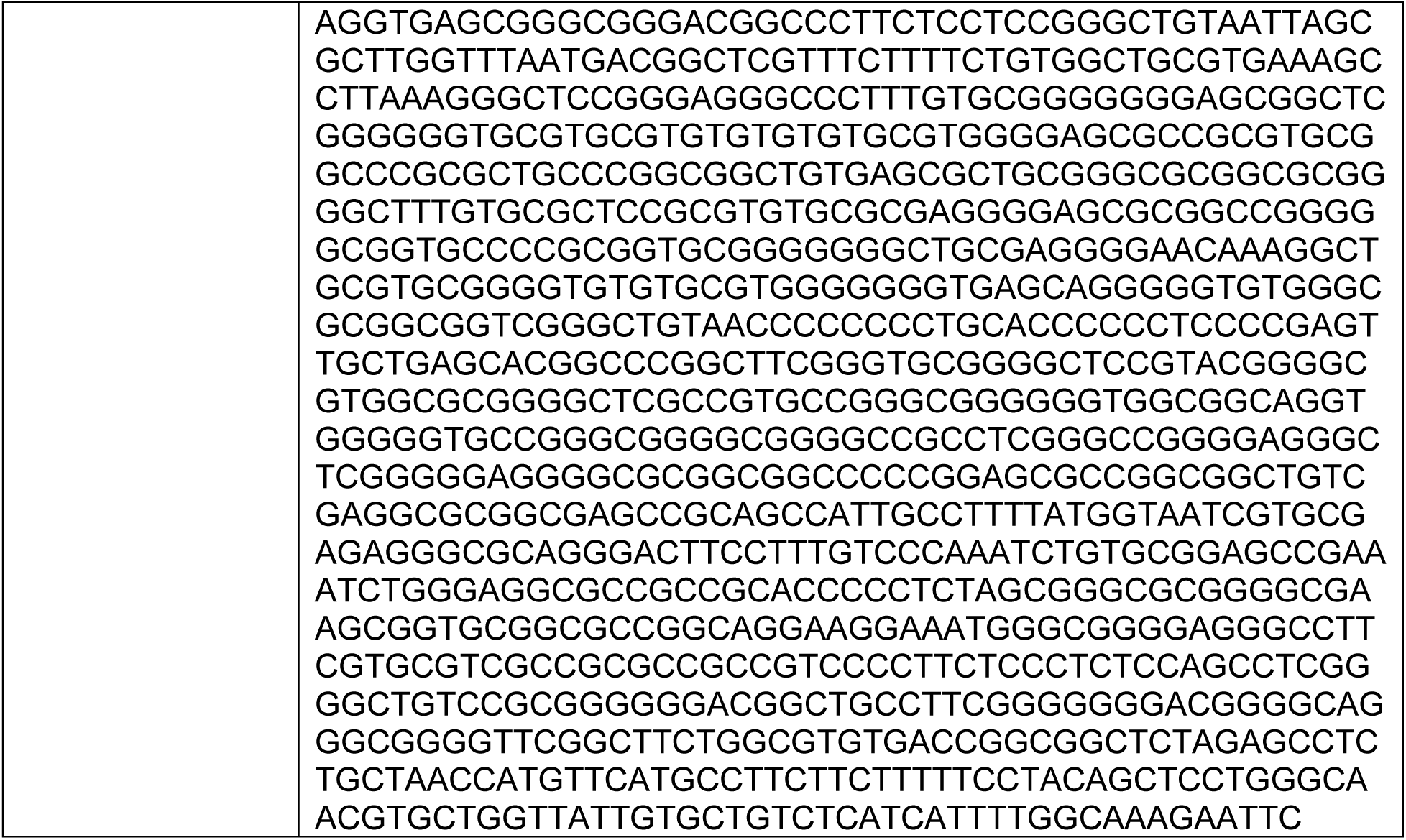
List of promoter parts used.

**Table S3:**
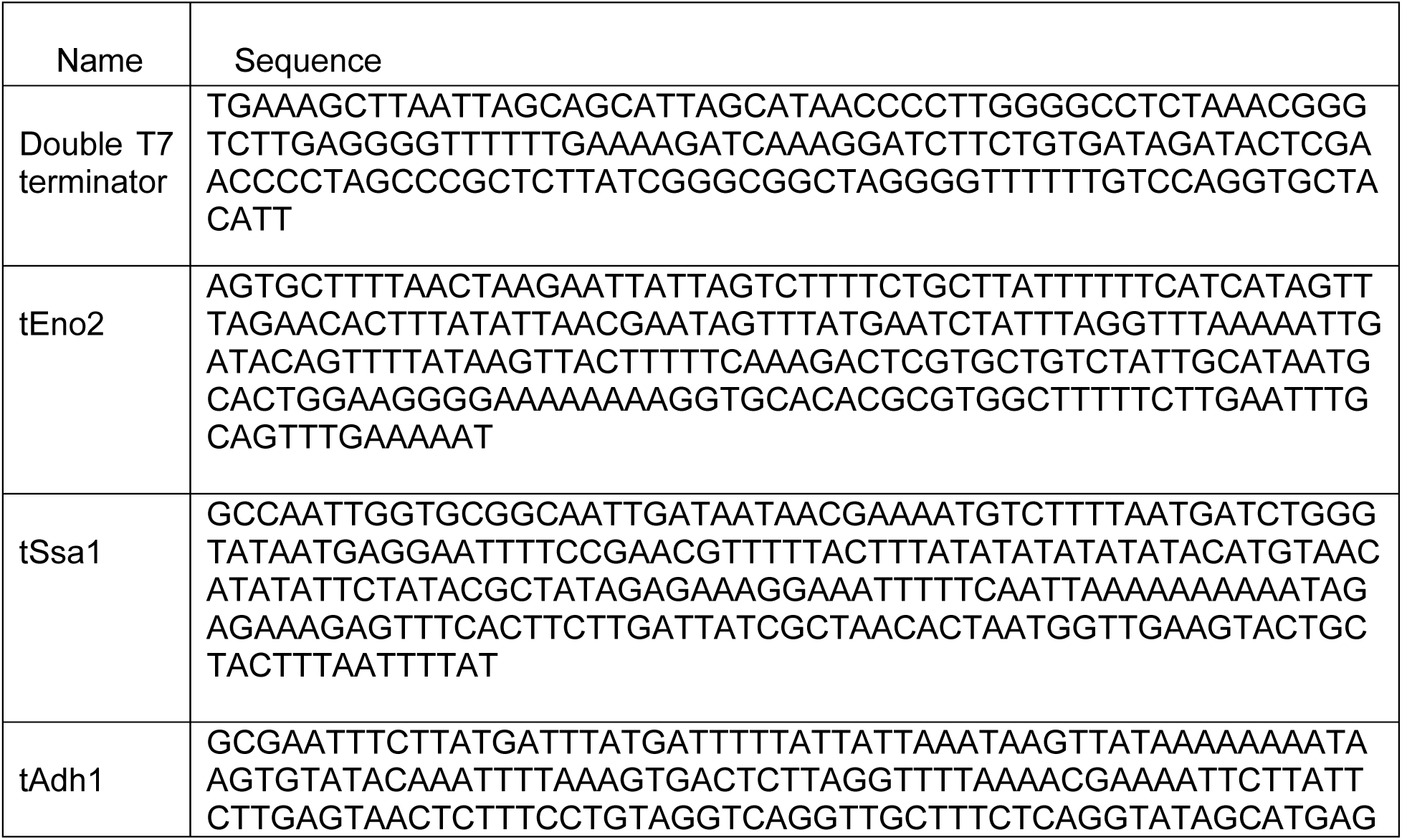
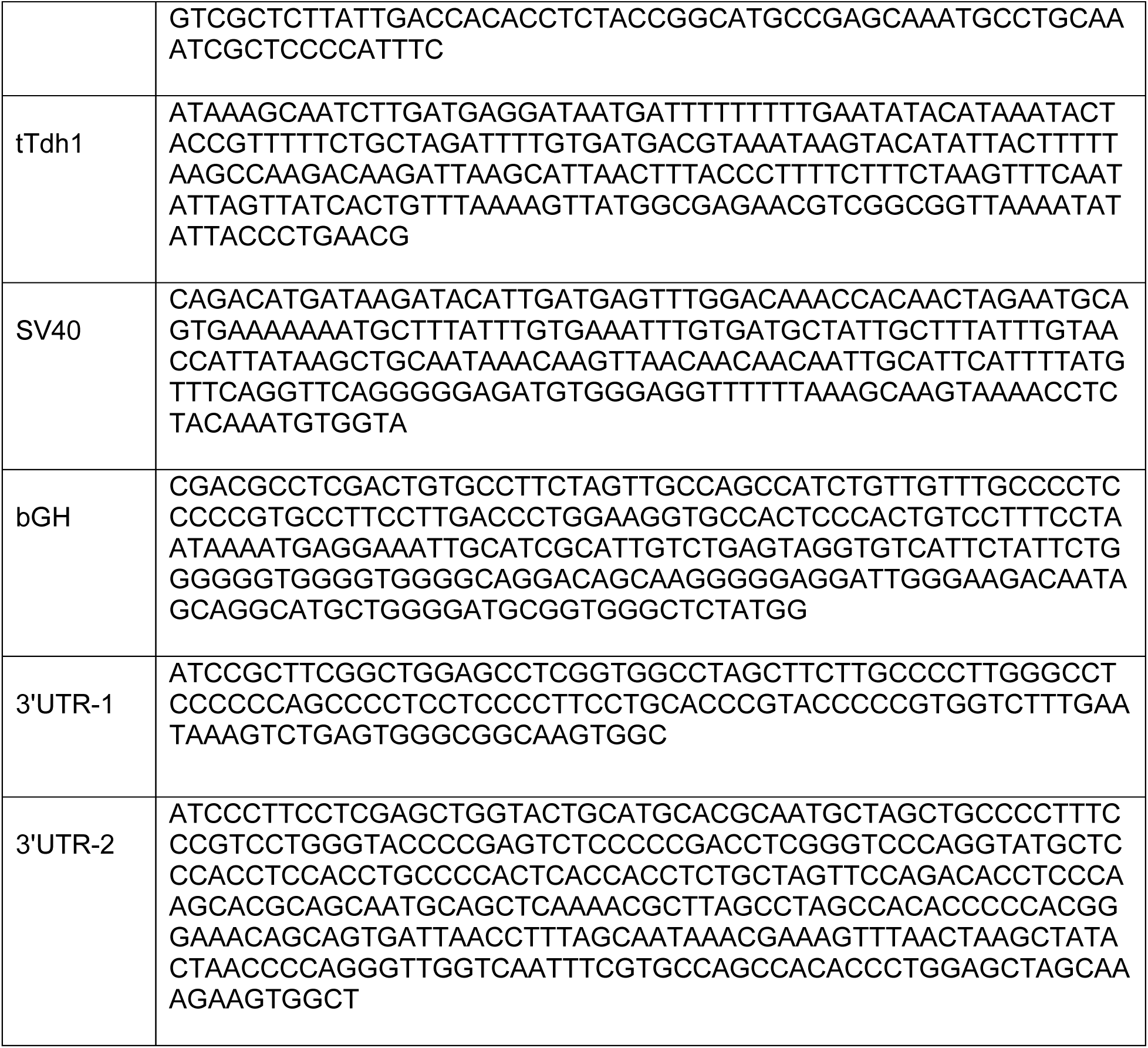
List of terminators, polyadenylation signals and 3’UTRs used.

**Table S4:** List of plasmid sequences included.

| Name | Description |
| --- | --- |
| ES205 | Target plasmid containing T7 promoter driving Zsgreen followed by a T7 terminator |
| ES246 | Expression vector containing wild type NP868R:T7 RNAP under Gal promoter |
| ES368 | Expression vector containing clone 368 under Gal promoter |
| ES369 | Expression vector containing clone 369 under Gal promoter |
| ES433 | Expression vector containing clone 433 under Gal promoter |
| ES443 | Expression vector containing clone 443 under Gal promoter |

## Materials and Methods

Cloning was performed from the MoClo yeast toolkit library and workflow. Parts not included in this library are described separately (**Table S1-3**). Plasmid assemblies were assembled from yeast toolkit parts and novel parts were ordered as gene blocks from IDT (Integrated DNA Technologies).

For the construction of each genetic element namely promoters, coding sequences and terminators, first they were checked for restriction sites for the following enzymes – BsmBI, BsaI, and NotI. The restriction sites in the coding sequences were removed by the use of synonymous codons while the other elements did not contain any of these restriction sites. The part plasmids were cloned into a common vector where each genetic element is flanked by Bsa1 restriction sites followed by appropriate overhangs. For the assembly of both single TU or multi-TU, the following procedure was used: 10 fmol of backbone plasmid and 20 fmol of parts/TUs were used in a 10uL reaction with 1ul of 10 × T4 ligase buffer along with 100 units of BsaI-v2 (single TU) or Esp3I (multi-TU or parts) and 100 units of T7 DNA ligase. The cycling protocol used is: 24 cycles of 3 min at 37 °C (for digestion) and 5 min at 16 °C (for ligation) followed by a final digestion step at 37 °C for 30 min and the enzymes were heat inactivated 80 °C for 20 min. All constructs were transformed into DH10B cells, grown at 37 °C using standard chemical transformation procedures. The colonies that lack fluorescence were inoculated and plasmids were extracted using Qiagen Miniprep kit according to the manufacturer’s instructions Plasmids were maintained as the following antibiotics kanamycin (50 μg/mL), chloramphenicol (34 μg/mL) and carbenicillin (100 μg/mL) wherever required. The plasmids were sequence verified by Sanger sequencing. The following enzymes were used for the assemblies – BsaI-v2 (NEB #R3733S), Esp3I (NEB #R0734S) and T7 DNA ligase (NEB #M0318S).

### Yeast transformations

Yeast were transformed with the Zymo EZ yeast transformation II kit (Zymo Research). For genomic integrations, 2 to 3 μg plasmids were linearized with NotI before transformation. 2 to 3 μg plasmid was digested with 1 μl NotI (NEB) for 1 h at 37 °C and heat inactivated prior to transformation. 5 μl of the digestion reaction was used for 100 μl cells.

### Yeast strains and functional assays

All strains were based on the BY4741 background (*S. cerevisiae*, S288C derivative) MATɑ his3Δ1 leu2Δ0 met15Δ0 ura3Δ0.

All engineered strains were grown overnight in SD at 30 °C for 20 h and their respective drop-out selective media dropout medium. Samples were diluted 1:10 in respective dropout media with 5% D-galactose (or varying levels as shown in the figures) and 2% D-raffinose as a carbon source. Samples were then grown at 30°C for 18 h. Cells were subsequently washed twice and resuspended in cold PBS. For cytometry, cells were diluted 1:100 in PBS, and assayed in SA3800 (Sony) spectral analyzer using autosampling. Samples were gated for singlets, and mean fluorescence was calculated for 10,000 events.

For sorting, library of the enzyme variants was constructed using error PCR and transformed into yeast strain containing the reporter construct. Following the induction, the cells were analyzed using the SH300 sorter (Sony). The cells were gated for singlets, and the top 1% ZsGreen expressing cells were sorted and recovered in the SD -His-Leu dropout media. Following their recovery, genomes were extracted and used for subsequent library generation. Following rounds of selection, specific colonies were isolated from the enriched populations by dilution plating, and their expression were characterized as described above. The best performing variants were sequenced to determine the mutations.

### Cell Culture and transfection

Human Embryonic Kidney 293 T cells (HEK293T, ATCC) were cultivated in High Glucose Dulbecco’s modified Eagle’s media (DMEM-high glucose, supplemented with 10% fetal bovine serum(FBS,GIBCO) All cell cultures were kept under 5% CO2 and 37°C temperature. All plasmids were transfected into cells by Lipofectamine 3000 (Invitrogen) according to the manufacturer’s instructions. Typically, 8×10^4^ cells were seeded on 24-well plates, and confluency of cells was around 70–80% on the day of transfection. Two days after transfection, the cells were trypsinized and collected in deep-well 96 well plates (Grenier). The collected cells were washed twice with PBSA (PBS + 1% BSA), and diluted 1:10 in 200uL prior to analyzing using the spectral analyzer SA3800 (Sony).

## Acknowledgements

ECG would like to thank the Genetic Design and Engineering Center (GDEC): A CPRIT Core Facility (RP210116) for supporting this work. ADE was supported by Welch F-1654; NIH R01-EB027202; and a Texas Biologics seed grant. JG acknowledges support from the Center for RNA Therapeutics, HMRI, the ADAPT fund, and a generous gift from the Jerold B. Katz Foundation.

## Author Contributions

SK and EG conceived of the study with input from JG. SK and EG performed the experiments with assistance from DB. All authors helped analyze the data. SK, JG wrote the manuscript with input from all authors. SK and EG led the study under the supervision of ADE and JG.

